# Bamboozling Interactions: Interspecific associations within mixed-species bird flocks in bamboo in the Eastern Himalaya

**DOI:** 10.1101/2023.10.08.561415

**Authors:** Sidharth Srinivasan, Aman Biswakarma, D.K. Pradhan, Shambu Rai, Umesh Srinivasan

**Affiliations:** Post-Graduate Programme in Wildlife Biology and Conservation, National Centre for Biological Sciences, Bangalore, India 560 065; Centre for Ecological Sciences, Indian Institute of Science, Bangalore, India 560 012

**Keywords:** arthropods, multi-species groups, network analysis, null model, resource availability, specialist species

## Abstract

1. Bamboo, although widespread, is a globally understudied habitat, but supports significant biodiversity, including specialist bird species. Mixed-species bird flocks (flocks) are an important and regular feature of forest bird communities worldwide and have been well-studied. However, how and why flocks might differ between bamboo and non-bamboo habitats is unknown.
2. We studied flocks and their arthropod prey in rainforest and bamboo stands across two seasons in the Eastern Himalaya. Using network analysis, we compared interspecific associations within flocks, and asked if resource availability (arthropod abundance) drives flocking.
3. Bamboo and rainforest flocks were significantly different, with bamboo flocks being more cohesive, less modular and their species more interconnected. Further, bamboo flocks were more consistent in their composition across seasons whereas rainforest flocks partially disintegrated in spring, probably because of increased arthropod abundance. In bamboo, the loss of arthropods in an entire substrate and the increase in other substrates was potentially insufficient to cease flocking in spring.
4. Resource availability and the unique foraging behaviour of the bamboo specialists likely drive flocking in the Eastern Himalaya bamboo. Our study is among the few to describe bamboo flocks and we emphasise conserving bamboo stands to ensure persistence of this unique flock in the Eastern Himalaya.

## INTRODUCTION

Numerous organisms, including mammals, fish and birds interact to form complex interspecific associations that structure communities and maintain community stability (Paijmans et al., 2019; Sridhar et al., 2009; Stensland et al., 2003; Thébault & Fontaine, 2010). Such associations are well-studied in insectivorous birds, in which an aggregation of two or more species, feeding together and moving in the same direction are known as mixed-species flocks (hereafter, flocks; (Kotagama & Goodale, 2004; Sridhar et al., 2009)). Many bird species participate in flocks, but some species are more important than others for flock initiation and cohesion; these are called ‘nuclear’ species. ‘Attendant’ species follow nuclear species. Associations between species can vary at the individual level and with species and environmental conditions and tend to be dynamic (Bangal et al., 2021; Goodale et al., 2017; Greenberg, 2000). Two main non-mutually exclusive hypotheses have been proposed to explain interspecific associations within flocks – improved foraging efficiency and lowered predation risk (Morse, 1977). Globally, a large body of literature has been devoted to investigating the structure, composition, and function of flocks (Sridhar et al., 2009, 2012).

Flocks are known to be structured by seasonality and the resulting fluxes in arthropod resources, especially in the Paleotropics and in subtropical and temperate forests. In highly seasonal areas, flocks are also seasonal (McClure, 1967); flocks in tropical regions, on the other hand, are considered to be stable and permanent throughout the year, especially in the Neotropics (Munn, 1985), and in a few Paleotropical systems (Jayarathna et al., 2013). Flock compositions and species’ roles in flocks also vary across time and habitats (Jayarathna et al., 2013; Jiang et al., 2020; Morse, 1970), and reduced food availability has been thought to drive mixed-species flocking, especially during periods of resource scarcity (U. Srinivasan & Quader, 2012). Resources in terms of arthropods show distinct seasonal fluctuations, and insectivorous birds are known to alter their ecology in response to these changes (Ghosh et al., 2011). However, almost all our understanding of flocks comes from forests in tropical and subtropical regions, and other habitats like bamboo have been widely neglected.

Bamboo (Poaceae: Bambuseae) is a distinct and diverse group of plants with a uniquely shared ecology. Overlapping with much of the range where avian mixed-species flocks occur, bamboo is found in tropical and subtropical habitats around the world, with Asia harbouring the maximum species richness, followed by South America (with less than half of Asia’s species richness; (Bystriakova et al., 2003, 2004)). However, our knowledge of birds in bamboo is heavily skewed towards the Neotropics, where over 90 species of primarily insectivorous Neotropical birds across different families are known to be partial to bamboo, with at least 20 species being obligate bamboo specialists (Areta et al., 2009; Kratter, 1997; Leite et al., 2013). Many species exhibit unique morphologies that allow for distinct foraging behaviours, such as the upcurved bill of the *Syndactyla ucayalae* (Peruvian Recurvebill), helping it split open bamboo stems to forage on arthropod prey. A recent study confirms the existence of specialist birds in bamboo with distinct foraging strategies in the Eastern Himalaya as well (S. Srinivasan et al., 2023)

Although some of these specialist species are also known to participate in flocks (Parker et al., 1997) and although studies on mixed flocks from the Neotropics are numerous, systematic studies on flocking assemblages in bamboo are limited not only from this region (Guimarães & Guilherme, 2021), but globally. Therefore, information about the structure and composition of flocks in bamboo is almost non-existent and patterns of interspecific associations between bamboo specialists/other species, and the mechanisms that drive flocking in bamboo remain unknown.

We used network analyses to investigate patterns of species associations in flocks in bamboo stands and rainforest in winter and spring in the Eastern Himalaya. Network theory has been increasingly used to study mixed-species systems as it helps quantify overall community structure and interspecific associations (Bangal et al., 2021; Montaño-Centellas, 2020; Montaño-Centellas et al., 2023; Proulx et al., 2005). Quantifying networks can help uncover network (flock)-level and species-level properties.

We asked the following questions and tested predictions related to flocks and resource availability in bamboo and rainforest in winter and spring in the Eastern Himalaya:

1. How does the structure and composition of flocks differ between bamboo and rainforest, across seasons? Given the presence of bamboo specialist species (S. Srinivasan et al., 2023), we predicted that flock networks and interspecific associations between species in bamboo would be stronger and more cohesive than flocks in the more diverse rainforest, especially in winter when resources might be limiting (Ghosh et al., 2011). We expected this based on similarities in behaviour in bamboo specialists that might confer foraging or anti-predator benefits through flocking (Goodale et al., 2020).
2. Do fluxes in resource availability drive the structure and composition of flocks, across both habitats and seasons? Given the inherent differences in arthropod availability in both habitats and seasons (S. Srinivasan et al., 2023), we expected the structure and the composition of flocks to be dissimilar between both habitats and seasons. With increases in arthropod abundance in spring, we expected flocking to reduce (and interspecific associations to weaken) in both habitats.

## METHODS

### Study Area

We carried out this study in the Eastern Himalaya, specifically the Eaglenest Wildlife Sanctuary (27.03° to 27.15° N and 92.30° to 92.58° E) in the West Kameng district in Arunachal Pradesh, India. Situated within the Himalayan Global Biodiversity Hotspot, this is a region of high bird and bamboo species richness (Bystriakova et al., 2003; Myers et al., 2000; Orme et al., 2005). Eaglenest Wildlife Sanctuary hosts a great diversity of habitats, from tropical rainforest to high-elevation coniferous and rhododendron forest across its elevational range.

We sampled bamboo and old-growth rainforest at altitudes ranging from 800-1200 m ASL in Eaglenest Wildlife Sanctuary. The vegetation in this area is tropical/sub-tropical wet evergreen forest (Champion & Seth, 1968) interspersed with primary, mature bamboo. In this region, bamboo tends to grow on steep slopes and the dominant bamboo species is *Dendrocalamus longispathus*, a large, clump-forming bamboo with sizable culms (stems) and relatively closed canopies, forming near-monotypic bamboo-dominated patches. The total area of the bamboo patches in the study area was ∼30 ha., with a few very large trees interspersed within these patches. We selected similarly sized undisturbed primary forest patches as the rainforest plots. The study period spanned two seasons, winter from January to mid-March 2022 and spring from mid-March to mid-May 2022.

### Sampling Design

#### Flock Sampling

We sampled flocks in line transects established in two habitats – rainforest and bamboo (seven transects in each habitat) and two seasons – spring and winter (see (S. Srinivasan et al., 2023) for a map of the transect locations). We also recorded flocks outside transects opportunistically, when walking between transects or on previously established trails in the two habitats. Because of the ruggedness of the terrain, adjacent transects were not likely to be independent of each other in terms of the home ranges of birds, especially in winter. Nevertheless, we treated each flock as an independent sample, because flocks provide a ‘snapshot’ of the realised interactions within them; members are dynamically participating in and exiting flocks within short time periods (Graves & Gotelli, 1993). We walked transects in the morning and evening but avoided very early mornings and late evenings when flocks would be forming or dissolving. We ensured that effort was similar across habitats in each season (47 vs 48.5 km in bamboo and rainforest in winter and 27 vs 24 km in bamboo and rainforest in spring).

We defined a flock as a group of at least two species, moving in the same direction (Kotagama & Goodale, 2004). We followed the ‘gambit of the group’ method, where individuals that were observed together during a sampling period were considered to be associated (Farine & Whitehead, 2015; Franks et al., 2010). On encountering a flock, we recorded the species composition of the flock by noting down the identity of the species, the group size of each species (i.e., the number of individuals) and the time. For flocks encountered outside transects, we also recorded the habitat in which they were observed. For gregarious species where accurate group size measurements were not always possible, we recorded a range, from the minimum to the maximum number of individuals in the group (Borah et al., 2018; U. Srinivasan et al., 2012) and we used the midpoint of this range in subsequent analysis. All the species recorded in a given habitat and season were pooled together to create a flock-by-species matrix.

#### Arthropod Sampling

To quantify resource availability, we sampled arthropods in both habitats and seasons. We selected 30 points at random in each habitat with a minimum distance of 30m between points. The points were randomly distributed around the transects, in each habitat and were sampled once in each season. We employed two commonly used methods – pitfall traps, for ground-dwelling arthropods (ground) and branch-beating, for foliage arthropods (foliage; (Cooper & Whitmore, 1990; Hohbein & Conway, 2018)). The bamboo specialist birds in this region were known to feed from within the bamboo sheaths (S. Srinivasan et al., 2023), and thus, we also sampled arthropods from this substrate (sheath; sampled only in winter, as the sheaths fall off the bamboo stems after winter), by adapting the branch-beating technique.

To capture ground-dwelling arthropods, we employed pitfall traps consisting of 5 cm diameter plastic containers buried flush with the soil. We filled these traps with a 10:90 detergent-to-water solution and left undisturbed for 48 hours. For foliage arthropods, we randomly selected branches 1-4 m in height and shook them into a large funnel-shaped beating bag for ∼10 seconds, allowing arthropods to be collected in a plastic container that was attached to its apex. To sample arthropods from bamboo sheaths, we held the beating bag close to the culms and dislodged them into the bag, shook it and collected the arthropods in the container at the end of the bag (S. Srinivasan et al., 2023).

We sampled a total of 270 points (150 in winter and 120 in spring), with each point being sampled in both winter and spring. We sampled during clear and sunny weather and stored the collected arthropods in 100% ethanol, which were later identified up to the order level.

### Analyses

#### Cluster Analysis

We used cluster analysis to categorise flocks into various types. Rainforest flocks from this region are known to be of three types – understory flocks, primarily comprising of small-bodied species, led by *Alcippe nipalensis* (Nepal Fulvetta), midstory/canopy flocks, led by *Melanochlora sultanea* (Sultan Tit) or by *Pericrocotus* sp. (minivets) and flocks of large-bodied species led by *Garrulax sp.* (laughingthrushes; (Borah et al., 2018; U. Srinivasan et al., 2012)). Moreover, it was apparent from fieldwork that there were at least two types of flocks in bamboo, one that was led by the *Gampsorhynchus rufulus* (White-hooded Babbler) and comprised the bamboo specialists, and the other led by *Alcippe nipalensis*. Therefore, we used a *k*-means cluster analysis to categorise flock types in each habitat and season and set the number of clusters to three. For further analyses, we then selected the clusters that contained the greatest number of flocks and with the presence of the *Alcippe nipalensis* for the rainforest flocks and the *Gampsorhynchus rufulus* for the bamboo flocks.

#### Structure and composition of flocks

To quantify differences in the composition of flocks, we first selected the flocks with the presence of *Alcippe nipalensis* in the rainforest flocks and *Gampsorhynchus rufulus* in the bamboo flocks in each season. We then constructed a flock-by-species matrix based on Hellinger-transformed raw counts of species (Buttigieg & Ramette, 2014). We employed the Bray-Curtis index on the transformed data (Bray & Curtis, 1957) and visualised the differences in flock composition by using a Non-metric Multi-Dimensional Scaling (NMDS). We used a PERMANOVA to test for significance (Permutational Analysis of Variance; (Anderson, 2017)).

#### Network Analyses

We used network theory to analyse the flocks and quantify interspecific associations in both habitats and seasons (Borah et al., 2018; Farine & Whitehead, 2015; Proulx et al., 2005). In these networks, nodes represent species and the edges between the nodes represent associations between species. After selecting flocks from the *k*-means cluster analysis, we constructed flock-by-species matrices and used these to construct weighted networks for each habitat and season, where the weights between species in the network represented the frequency of distinct co-occurrences of species. For every network, we calculated network- and node-level metrics.

At the network level, we calculated network density and modularity. These metrics quantify the overall connectivity (density) and cohesion (modularity) of a network (Montaño-Centellas, 2020). Network density is the proportion of all possible connections in a network that are observed (Newman, 2003). Network density, therefore, represents distinct co-occurrences or associations between species in flocks and can help understand the overall connectedness of the flock. Higher values indicate a denser, highly connected network. Modularity calculates the strength of different modules/clusters (closely connected nodes) in a network by estimating how well-separated modules in a network are based on the number of edges within and across modules (Clauset et al., 2004). We calculated modularity using the Louvain method with a community detection algorithm (Bangal et al., 2021). Modularity represents the overall cohesion of the network – higher values indicate when these subsets of species interact more strongly within them and weakly with the other species in the network.

We also calculated node-level (species-level) metrics which quantify the importance of a node within a network. We calculated degree, which represents the number of edges connected to a particular node, and weighted degree (association strength), which is the sum of the frequency of distinct co-occurrences for each node. These measures are sensitive to species richness; therefore, we normalised degree by dividing values by the number of possible connections in a flock (*n* – 1, where *n* is the total number of species recorded participating in the habitat- and season-specific flock) and weighted degree by dividing values by the maximum value of weighted degree in each network. Degree represents the interspecific associations of each species while weighted degree is a measure of centrality, which captures the structural importance of a species in a network. We also visualised the distribution of degrees of the weighted networks, to understand the structure of the networks across habitats and seasons. Supplementary Material Table S1 summarises the network definitions and their biological interpretations.

#### Random networks

We simulated null networks based on the observed data for each habitat and season to test whether network- and species-level metrics deviated significantly from random expectations. We randomised species’ participation within flocks while controlling for flock species richness and the number of flocks that a species participated in each network (i.e., randomising matrix entries while holding row and column totals constant; 27,45). Thus, the null model preserves differences in species richness among habitats and seasons but assumes that differences in flock participation within each habitat and season are proportional to observed species occurrences. We simulated 10,000 null flocks and compared the distribution of the null metrics with the metrics calculated from the selected networks from the *k*-means cluster analysis (‘observed flocks’) to infer if the values lie within the null expectation (‘null flocks’). This iterative permutation procedure allows us to test if interspecific associations are potentially caused by underlying social interactions between individuals or by some other factors, such as congregations between individuals due to their specificity for habitat or resource (Farine, 2017; Montaño-Centellas, 2020).

#### Arthropod abundances

We analysed per-point arthropod abundance in both habitats and seasons, across substrates. The order Hymenoptera (mostly ants) was removed before analysis, as it increased considerably in the spring, making it harder to compare abundances. We encountered a single point in bamboo pitfall traps in the spring that contained a hyper-abundance of Isoptera (875 individuals). This was clearly an outlier and therefore, we removed Isoptera as well.

We carried out all analyses in program R (R Core Team, 2022). We created random networks using the package *EcoSimR* (Gotelli et al., 2015) and network analyses using the package *igraph* (Csardi & Nepusz, 2006).

## RESULTS

We observed 2,004 individuals represented by 96 species in 304 flocks across both habitats and seasons. We filtered these to remove rare species (occurring in less than 1% of all flocks) and categorised them by habitat and season.

### Cluster Analysis

The *k*-means cluster analysis categorised the flocks in each habitat and each season into three types. We selected the type with the most number of flocks and with the presence of the *Alcippe nipalensis* in rainforest flocks and *Gampsorhynchus rufulus* in the bamboo flocks. Details of the number of flocks selected for each habitat and season based on the cluster analysis are presented in (Table 1).

**Table 1.**
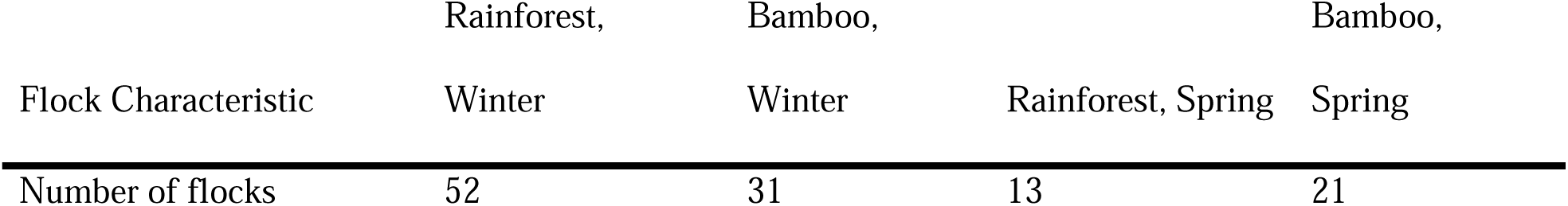
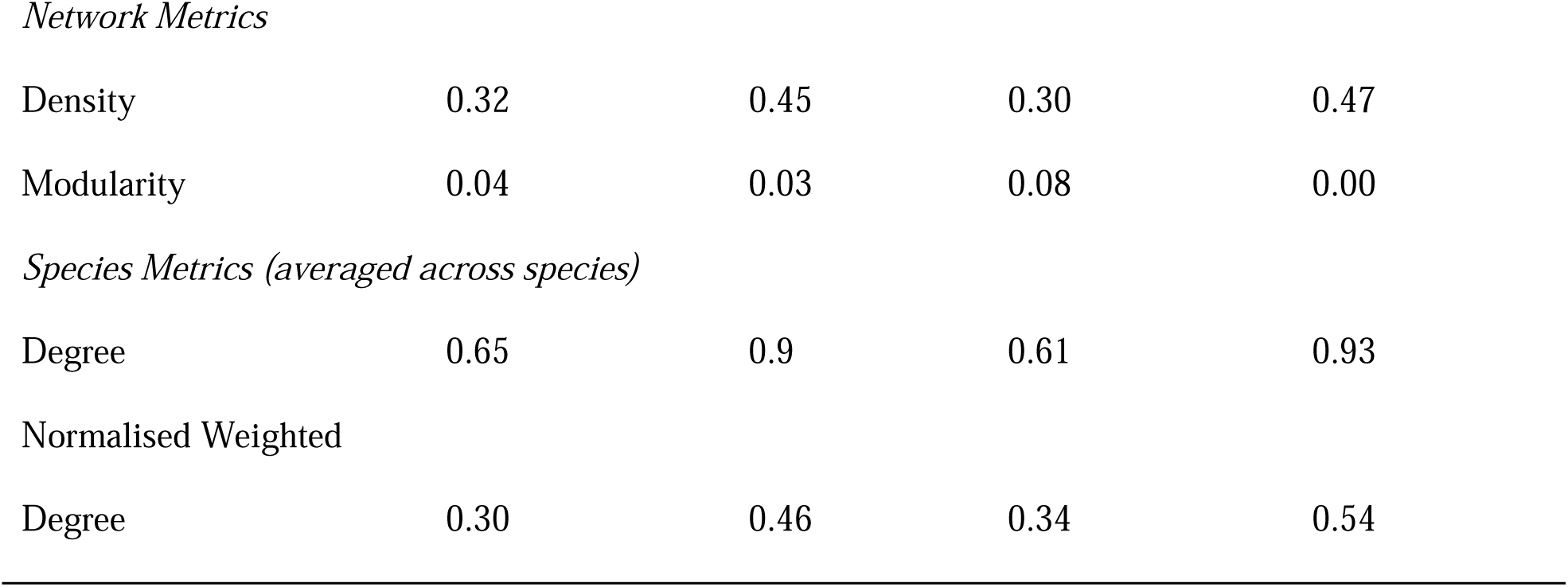
- Characteristics of flocks used in the analyses. Number of flocks represents the total number of flocks selected in each habitat and season based on the *k*-means cluster analysis. For definitions and biological interpretations of the metrics, please refer to Supplementary Material Table S1.

### Structure and composition of flocks

The NMDS separated the flocks by habitat on the first axis well and indicated the presence of a significantly different flock in each habitat (stress = 0.08, *R^2^* = 0.5, *F*_1,138_ = 139.78, *p* < 0.01; Figure 1). Flocks were composed of species that were distinct in each habitat and few species were common across all habitats and seasons. Rainforest flocks were mostly composed of *Alcippe nipalensis*, *Dicrurus remifer* (Lesser Racket-tailed Drongo), *Phylloscopus castaniceps* (Chestnut-crowned Warbler), and *Culicicapa ceylonensis* (Grey-headed Canary Flycatcher) in winter, while in the spring, *Phylloscopus reguloides* (Blyth’s Leaf Warbler) and *Phylloscopus whistleri* (Whistler’s Warbler) replaced the latter two winter species as the most common species. In bamboo, the most common species were *Dicrurus remifer*, *Gampsorhynchus rufulus*, *Alcippe nipalensis*, *Phylloscopus poliogenys* (Grey-cheeked Warbler), and the *Suthora atrosuperciliaris* (Pale-billed Parrotbill), in winter; in spring, *Gampsorhynchus rufulus*, *Suthora atrosuperciliaris* and *Alcippe nipalensis* were encountered the most. The composition of flocks in bamboo differed from that of the rainforest mainly due to the presence of bamboo specialist species such as *Gecinulus grantia* (Pale-headed Woodpecker), *Pomatorhinus ochraceiceps* (Red-billed Scimitar Babbler), *Abroscopus superciliaris* (Yellow-bellied Warbler), *Gampsorhynchus rufulus* and *Suthora atrosuperciliaris*, which were entirely absent from rainforest flocks (S. Srinivasan et al., 2023). Likewise, the rainforest flocks harboured a few species that were never encountered in the bamboo flocks; for instance, *Parus monticolus* (Green-backed Tit), *Mixornis gularis* (Pin-striped Tit-Babbler) and *Zosterops palpebrosus* (Indian White-Eye). Seasonally, flocks were not dissimilar, however, there were slight differences due to the absence of the winter migrants in the spring, as migrants moved to breed at higher elevations. This difference was marked in the bamboo flock in spring, as it was composed entirely of the bamboo specialists, along with *Dicrurus remifer*.

**Figure 1.**
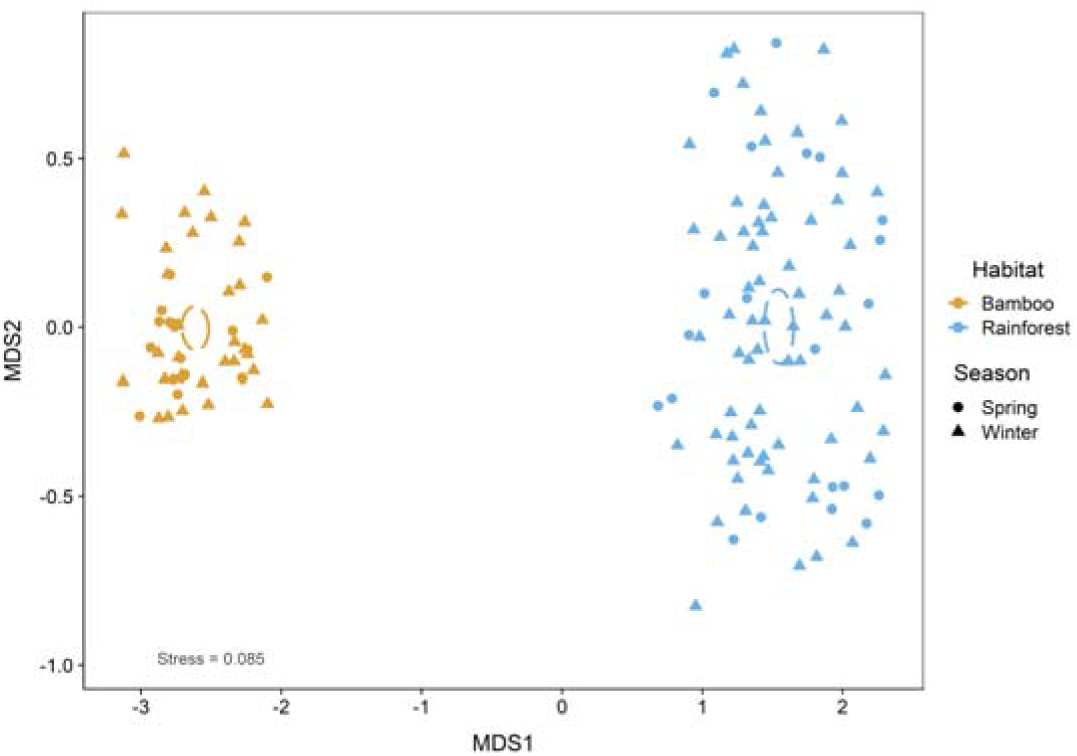
- Non-metric Multi-Dimensional Scaling representing dissimilarity between winter and spring flocks in bamboo and rainforest. Each point represents a flock in a season and habitat. Ellipsoids are centred around the mean and represent 95% confidence intervals around the points.

### Network Analyses

Network structure from the observed flocks was distinct across habitats and seasons (Figures 2, 3, Table 1). Network-level metrics from the observed networks indicated that bamboo flocks were more connected (Figure 2**A**), less modular (Figure 2**B**), and had more (Figure 3**A**) and stronger interspecific associations (Figure 3**B**), irrespective of the season. Bamboo flocks were denser than rainforest flocks, implying that there were more realised interspecific connections in the flock out of all possible connections. Across seasons, bamboo flocks were less modular, suggesting a more cohesive flock that had fewer divisions and more connections among adjacent species than in rainforest. Normalised degree, a species-level metric, also indicated that species in the bamboo flocks were more connected across seasons. The normalised weighted degree or association strength was higher in bamboo flocks and increased as the season progressed from winter to spring in both habitats (Figure 3**B**, Table 1). With consistently higher values for density and lower values for modularity, bamboo flocks appeared to be more structured than rainforest flocks, across seasons. Rainforest flocks were less connected and less cohesive than bamboo flocks. Across seasons, rainforest flocks became more sparsely connected, with values for density and normalised degree decreasing from winter to spring (Figures 2, 3). Rainforest flocks were also more clustered, especially in spring. Surprisingly, the normalised species’ association strength increased from winter to spring in this habitat, but only slightly (Figure 3**B**). The metrics for observed flocks differed significantly from the results of the randomised networks (Figures 1, 2). The exceptions were the values for modularity in the winter rainforest flocks and weighted degree in the winter bamboo flocks, albeit marginally.

**Figure 2.**
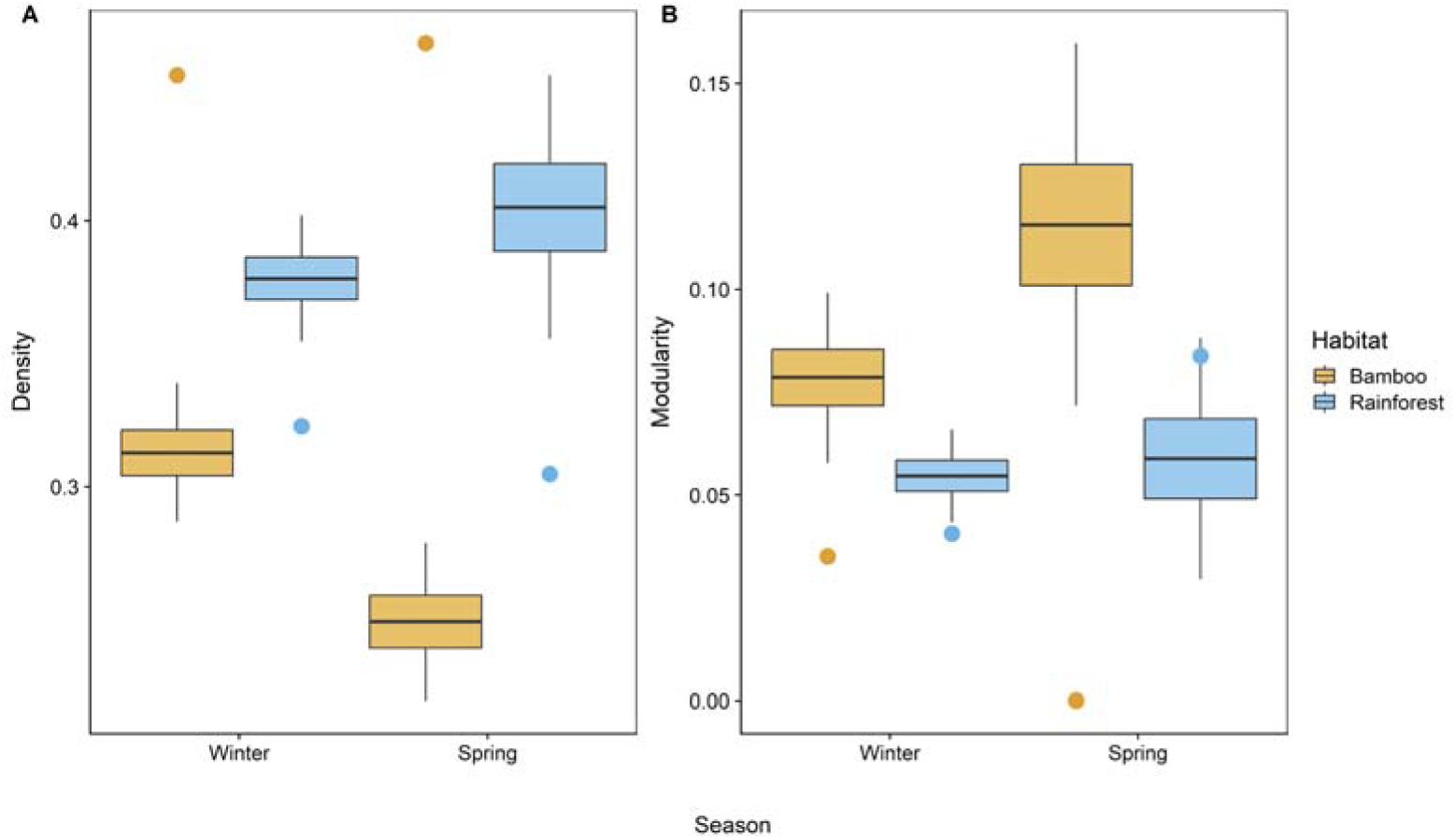
- Network-level metrics for flocks in both habitats and seasons. Network-level metrics represent density (**A**) and modularity (**B**) Dots represent values of each metric from the observed flocks while the box plots depict the mean and the standard deviation and the whiskers represent the minimum and maximum of the metrics from the null models.

**Figure 3.**
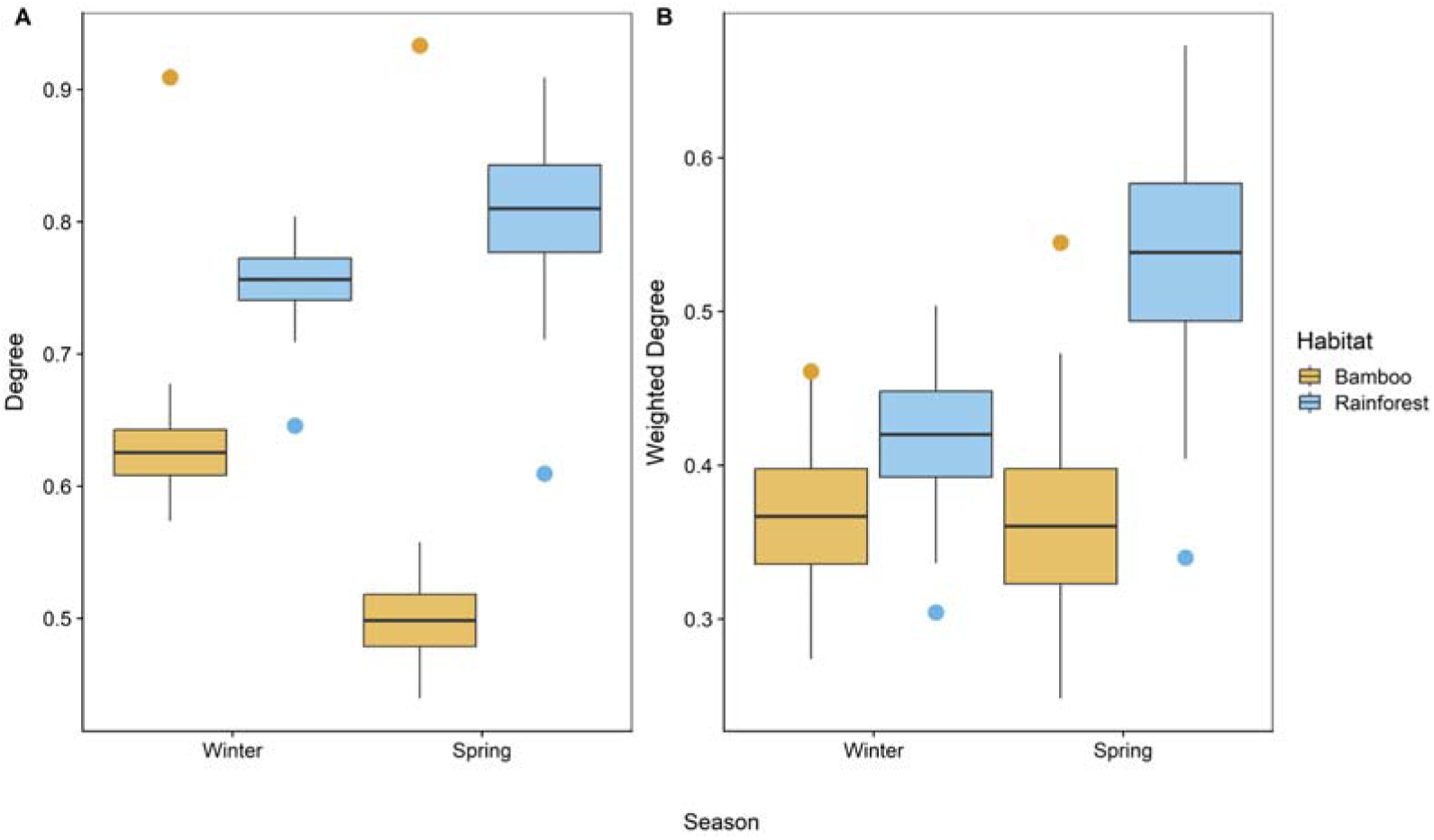
- Species-level metrics degree (**A**) and weighted degree (**B**) for flocks in both habitats and seasons. Dots represent values of each metric from the observed flocks while the box plots depict the mean and the standard deviation and the whiskers represent the minimum and maximum of the metrics from the null models.

The normalised frequency distribution of interspecific interactions (association strength) for each habitat and season indicated that rainforest had very few species that were highly connected with each other with most species only sparsely associated (Supplementary Material Figure S2). This was especially true in winter, where only a handful of species had very high weighted degree values and most species had much lower values, indicating a skewed network with sparse connections. In spring, the steepness of the curve seemed greater, with even fewer species being very highly associated with others and most species having few connections with others. Thus, relative to the total number of species, more species seemed to have higher weighted degree values in bamboo than in rainforest, indicating stronger associations in bamboo.

### Arthropod abundances

Sampling across habitats and seasons yielded 7,354 individuals representing 21 orders. The major orders were Araneae, Hymenoptera (> 90% were ants), Acarina, Diptera, Blattodea, Hemiptera, Orthoptera, Psocoptera, Coleoptera and Collembola. Arthropod abundances increased greatly in spring across substrates, irrespective of the habitat. The abundances in both habitats seemed to be comparable in winter (notches overlapped considerably; Figure 4) but the abundance of arthropods in rainforest was distinctly higher than that of bamboo in the spring. This was apparent in both the ground and the foliage-dwelling arthropods. The abundance of arthropods in sheaths in bamboo was not considerably different from their abundances in other substrates within bamboo or rainforest, in winter. In spring, however, the sheaths fall off the bamboo stems and the arthropods are unavailable in this substrate.

**Figure 4.**
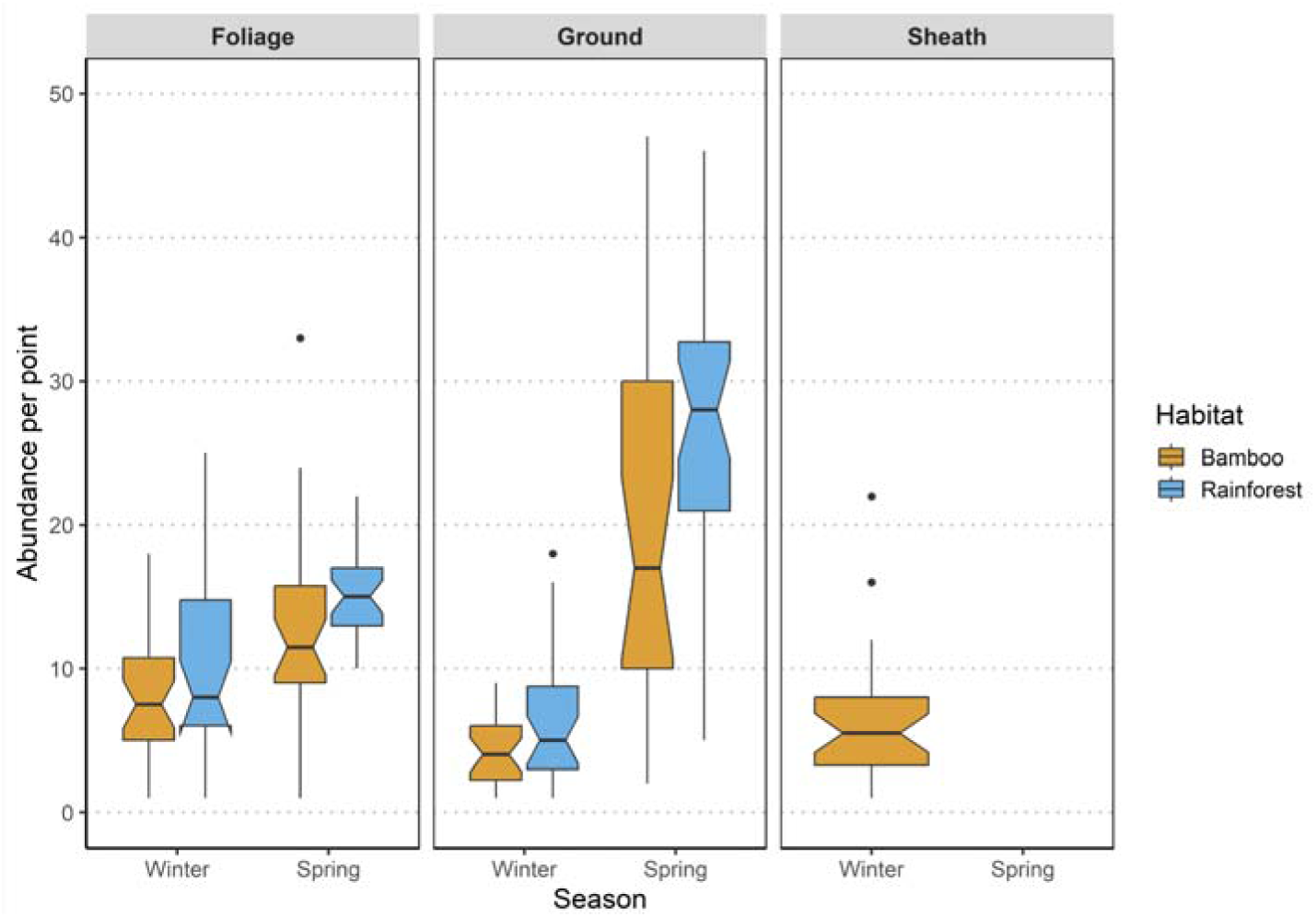
- Notched box plot depicting per point arthropod abundances by substrate, in each season and habitat.

## DISCUSSION

We provide one of the first quantitative assessments of mixed-species bird flocks in bamboo from anywhere in the world. We found that in the Eastern Himalaya, the structure and composition of bamboo flocks are distinct from those in rainforest, across seasons (Figures 1, 2, 3). Bamboo flocks mainly consisted of bamboo specialist species, and while these species readily associated with each other, they rarely associated with species that were not bamboo specialists. The results of the cluster analysis separated this flock from the other flocks in bamboo, corroborating observations from the field.

### Flock structure and composition

Flocks were composed distinctly between habitats, with flocks from bamboo separating well from those in rainforest (Figure 1). This indicates the uniqueness of the bamboo flock in the Eastern Himalaya, which is mostly composed of species that are bamboo specialists (S. Srinivasan et al., 2023). These specialist species associated with each other regularly in the bamboo flocks across seasons, making this flock a bamboo specialist species flock. *Gampsorhynchus rufulus* possibly acts as the ‘leader’ or the ‘nuclear’ species in these flocks, responsible for the initiation and/or maintenance of flocks. Corroborating field observations, *Gampsorhynchus rufulus* had the highest association strength/weighted degree (1.00) across both seasons, and was very vocal and interspecifically gregarious, making it an ideal nuclear species (Goodale et al., 2015; Goodale & Beauchamp, 2010; U. Srinivasan et al., 2010). Frequent non-bamboo specialist species that were found in bamboo flocks in winter include *Dicrurus sp.* (drongos) and these species are known to be excellent ‘sentinels’ (Goodale & Kotagama, 2005; Sridhar & Shanker, 2014). Sentinel species such as drongos are salliers that forage in flocks to feed on prey that is usually flushed out by gleaners (Satischandra et al., 2007; U. Srinivasan & Quader, 2012). These species are also very vocally active, providing reliable social signals to other species in the flock in the form of anti-predator alarm calls (Goodale & Kotagama, 2005). Therefore, drongos in the Eastern Himalayan bamboo flocks might be acting as sentinel species, salling and foraging on the prey flushed out by the bamboo specialists. Attendant species found in bamboo flocks in winter such as the *Cissa chinensis* (Common Green Magpie; 0.22) and the *Lalage melaschistos* (Black-winged Cuckooshrike; 0.25) were probably opportunistic flocking species in bamboo.

On the other hand, rainforest flocks were not dissimilar from flocks commonly encountered in forests from other studies in this region (U. Srinivasan et al., 2012). *Alcippe nipalensis* was possibly the nuclear species in this habitat (U. Srinivasan et al., 2012; Zhou et al., 2019), with high association strengths across seasons (0.92 in winter, 1.00 in spring). Other species that had relatively high association strengths include *Phylloscopus castaniceps* (1.00 in winter) and *Culicicapa ceylonensis* (0.81 in winter), which are common small-bodied understory attendant species in forest flocks in this region (U. Srinivasan et al., 2012). *Dicrurus remifer*, likely acting as the sentinel species in this habitat as well, also had relatively high association strengths, especially in winter (0.91 in winter, 0.45 in spring).

These findings were corroborated by other metrics from the network analyses, which revealed significant differences in flock structure between both habitats and seasons. Bamboo flocks were denser (Figure 2**A**) and more cohesive than rainforest flocks (Figure 2**B**, Table 1). Moreover, species in bamboo flocks had more (and stronger) interspecific connections (Figure 3). Contrarily, rainforest flocks had fewer and weaker interspecific associations and harboured more clusters within flocks, especially in spring. As the season progressed from winter to spring, rainforest flocks became more loosely associated, had fewer connections and were highly modular, signalling the (at least partial) breakdown of flocks in spring. Overall, the structure of the flocks in bamboo were tighter and had more consistent flocking across seasons, indicating the distinctiveness of the bamboo flock when compared to the rainforest flock.

The observed values of nearly all network and species-level metrics differed significantly from null expectations, the exception being the values for modularity in the rainforest flocks in winter and weighted degree in the bamboo flocks in winter, albeit marginally (Figures 2, 3). The significant departure from null indicates that the flocks are structured by interspecific or social interactions rather than random aggregations of individuals because of the habitat or underlying resources. While this was expected for the flocks in rainforest, this finding is significant for flocks in bamboo. Because the bamboo flock is almost entirely composed of bamboo specialist species, their associations could be interpreted as random aggregations because of the habitat. Therefore, the results from the null models point to underlying social interactions within flocks that drive flocking, especially in bamboo in the Eastern Himalaya.

### Seasonal arthropod availability might drive flocking

The Eastern Himalaya is a monsoonal system with spring and winter representing distinct wet and dry periods respectively but seems to host both seasonal and permanent flocks. Although both bamboo and rainforest flocking decreased with the onset of spring, this decrease was far greater in rainforest (Supplementary Material Figure S1). With resident birds occupied with breeding activities in the spring, this decrease in flocking in both rainforest and bamboo was not unexpected. However, rainforest flocks partially disintegrated in spring, as evidenced by the network- and species-level metrics (Figures 2, 3). This breakdown of flocking in rainforest was probably due to the increase in arthropod availability in spring (Figure 4). However, contrary to our expectations, the bamboo specialist flock system did not break down in spring, even though arthropod availability in the habitat increased (Figure 4). This increase was greater in rainforest in spring, and arthropod abundances across substrates were significantly higher when compared to bamboo (Figure 4). While this might have been sufficient to contribute to the partial breakdown of flocking in rainforest (U. Srinivasan & Quader, 2012), similar increases in arthropod abundances in other strata of bamboo were probably not sufficient to compensate for the loss of sheath arthropods. This could explain why bamboo flocks do not disintegrate during the spring and rainforest flocks do (at least partially), as arthropod resources in bamboo might not be as abundant as they are in rainforest (Figure 4). Bamboo flocks, composed of specialist species, seem to be obligate in bamboo, and resource availability might be a factor in driving flocking throughout the year.

### Benefits and costs associated with flocking in bamboo

Individuals join flocks based on the risks and benefits associated with flocking. (Sridhar et al., 2009; Sridhar & Shanker, 2014). Associating in flocks can help individuals spend more time foraging and less time being vigilant (Sridhar et al., 2009) and potentially enhance the fitness of individuals, possibly because of the benefits accrued (Dolby & Grubb jr, 1998; U. Srinivasan, 2019). Species can provide complementary (benefits provided by a flocking individual that are lacking in its associate) or supplementary benefits (benefits provided by a flocking member that are similar to its associate) in flocks with respect to anti-predator and foraging benefits (Goodale et al., 2020).

In the Eastern Himalayan bamboo flock, supplementary benefits could be provided by the bamboo specialist species. Because they employ the same foraging strategy (S. Srinivasan et al., 2023), social copying - where one individual observes and copies the foraging location or some other information related to foraging - is likely (Krebs, 1973). Birds are known to visit locations where their associates have successfully foraged (Waite & Grubb Jr, 1988). Members of the flock might also have information on previous foraging sites and might help participants avoid sites that have already been exploited (foraging efficiency theory, Diamond, 1981). In addition, complementary benefits can also be gained in tandem. The bamboo specialists might have traded off their vigilance for their specialised feeding behaviour, peering into the sheaths, internodes, and holes in bamboo or tearing apart the sheaths (Jones et al., 2020). Hence, these species might require others to alert them about the presence of a nearby predator. The sentinel species usually carry out this role but it can also be taken up by the nuclear species – drongos and *Gampsorhynchus rufulus* respectively in this case. The latter is known to forage in family groups, with juveniles and adults participating in flocks. Hence, kin selection would predict that *Gampsorhynchus rufulus* would warn their kin in the presence of a predator (Hamilton, 1964), and it would not be surprising to learn that other bamboo specialist species, such as *Suthora atrosuperciliaris* might have evolved to eavesdrop on these calls. Moreover, juveniles of this species have rufous and not white heads, closely resembling the parrotbills (Zhou et al., 2019). This could help them appear inconspicuous within the flock and survive to adulthood via predator dilution or other benefits gained through mimicking the more dominant, nuclear species (Zhou et al., 2019). Similarity in traits such as diet, body size and foraging behaviour might be key to acquiring these benefits (Goodale et al., 2020; Sridhar et al., 2009, 2012; Zhou et al., 2019). These complementary and supplementary benefits could be influenced by resource availability – when sheath arthropods are present in winter, individuals in bamboo flocks might rely on the complementary benefits provided by the drongos to alert them about a potential predator, while they peer/tear up the sheaths to find arthropod prey. However, the benefits acquired by these species in spring might require research, as overall arthropod numbers increase and sheath arthropods are lost.

There are some costs associated with flocking together as well - the more similar the niches of species in a flock, the more they might compete for the same resources. This presents a potential trade-off to bamboo specialists in flocks; however, such competition might still be lower than if they were in purely intraspecific flocks. Studies suggest that by not having to ‘match their activity’ (i.e., change behaviour to associate together in flocks), they might incur lower costs and find flocking more beneficial than intraspecific flocking (Sridhar & Guttal, 2018). Thus, the potential benefits of more efficient foraging might outweigh the competition costs of associating within flocks in bamboo.

### Future directions

Bamboo is unique in its ecology because of its boom-and-bust cycle, where both habitat and food resources are known to drastically fluctuate over its lifetime (Gadgil & Prasad, 1984; Janzen, 1976). During die-off events when arthropod resources are likely scarce, the bamboo insectivore specialists disperse to nearby bamboo stands (Areta & Cockle, 2012). Associating with interspecifics could be a strategy to help find food or even alternate bamboo patches during this period. Because the species in Eastern Himalayan bamboo flock have high association strengths (Figures 3**B**, Supplementary Material Figure S2), we could expect this to be the case during die-off events in this region. Investigating the fate of this flock during die-off events would be a key area of research.

Bamboo in this region is also an important resource to resident human communities and is harvested widely (Bystriakova et al., 2003). Thus, bamboo specialist species are forced to cope with an even more variable and fluctuating habitat, due to anthropogenic use. This could potentially threaten specialist species that seem to be highly interdependent, reliant on flocking in bamboo and are exclusive to this habitat. Additionally, bamboo specialists and bamboo flocks are known to have very small home ranges (Guimarães & Guilherme, 2021; Lebbin, 2013) and given their interdependency, individuals of the Eastern Himalayan bamboo specialist species likely exhibit flock fidelity. This could potentially affect the fitness of other species in the flock, depending on their importance within the flock, because the loss of nuclear species in flocks is known to reduce the fitness of other participant species (Dolby & Grubb jr, 1998). Bamboo flocks could potentially be less resilient to disturbances, because of the short tail in the weighted degree distribution (Supplementary Material Figure S2; (Thébault & Fontaine, 2010)). Therefore, investigating the responses of this flock to anthropogenic disturbances is a potentially important research area. Nevertheless, we emphasise the need to conserve large, contiguous bamboo stands at the landscape level in the Eastern Himalaya, which might be key to the survival of this highly specialised flock.

## CONCLUSIONS

Our study is one of the first to quantitatively describe mixed-species bird flocks in bamboo globally. Flocks in bamboo and rainforest in the Eastern Himalaya showed considerable differences in their structure and composition across two seasons. Bamboo flocks were more highly connected, had stronger interspecific connections and were less modular than the rainforest flocks in both winter and spring. Bamboo flocks were mainly composed of only bamboo specialists and were also more consistent across seasons, while the flocks in rainforest were more diverse and highly variable. Flocking seemed to be largely driven by the seasonal availability of arthropod resources in both habitats. The unique foraging behaviour of bamboo specialists, and the relative costs and benefits of associating in flocks can also help discern the patterns of grouping that are seen in the Eastern Himalayan bamboo flocks. Conservation of bamboo stands in the Eastern Himalaya might thus be essential for the persistence of these obligate bamboo specialist flocks.

## ACKNOWLEDGEMENTS

We thank the National Centre for Biological Sciences-Tata Institute of Fundamental Research (NCBS-TIFR) and the Nature Conservation Foundation for institutional and administrative support. We are thankful to the Arunachal Pradesh Forest Department for issuing permits to carry out this study (permit no. CWL/GEN/2018-19/Pt.IX/NG/3480-81). SS is grateful to The Habitats Trust for providing him with a grant to carry out this study. The Indian Institute of Science provided additional financial support. Rinchen Angmo and Yeswanth H.M. helped greatly with arthropod identification and Binod Borah provided useful discussions. We also thank Mangal Rai, Nima Tamang, S.K. Pradhan, Bharat Tamang and Binod Munda for their support and help during fieldwork.

