## Supplementary Material for "Bamboozling Interactions: Interspecific associations within mixed-species bird flocks in bamboo in the Eastern Himalaya"

**FIGURES**

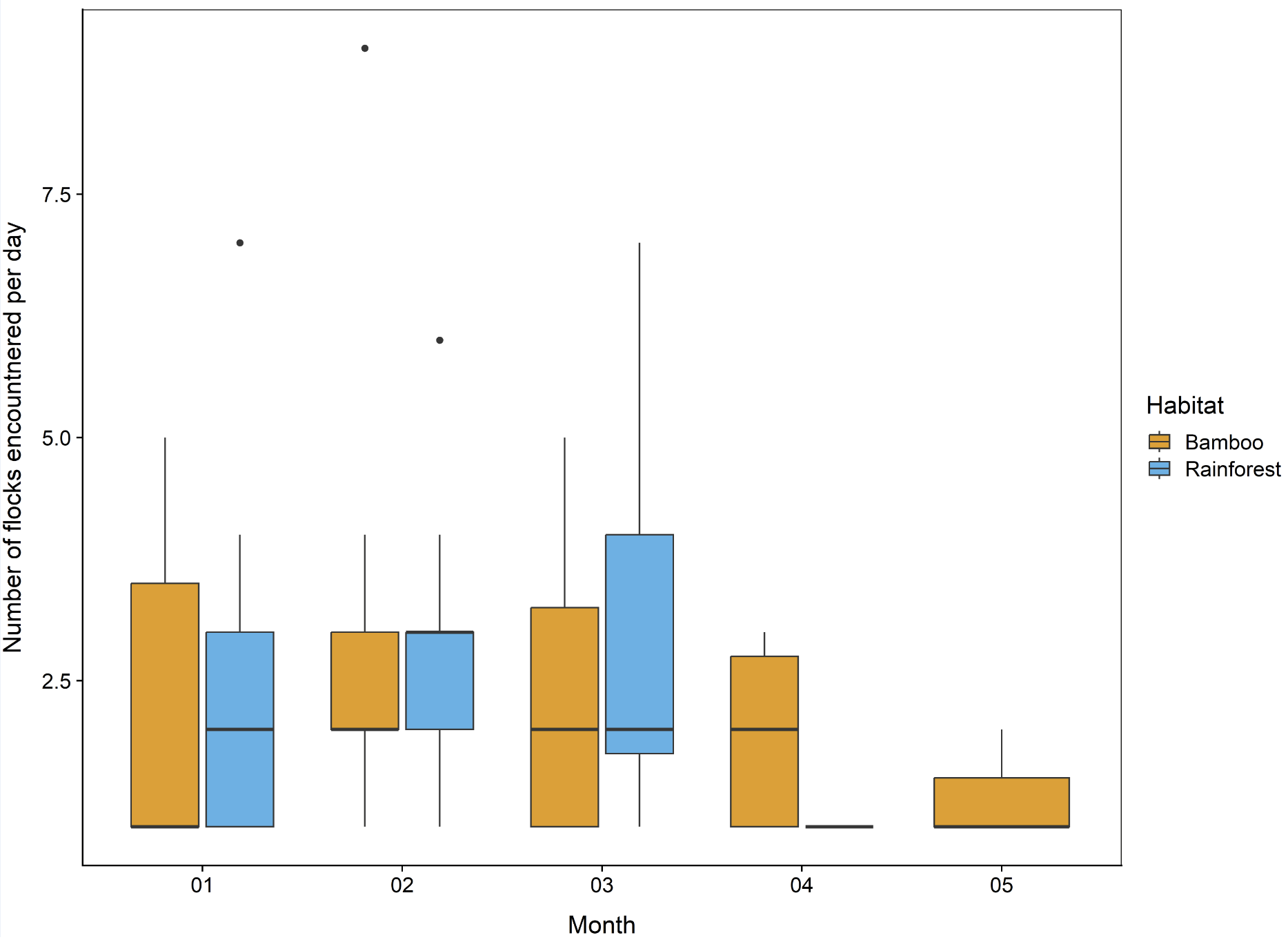

**Figure S1** - Number of flocks encountered per day, across months. January to mid-March is defined as winter and mid-March to mid-May is defined as spring. Whiskers in the plot represent 1.5 times the interquartile range and the points are outliers.

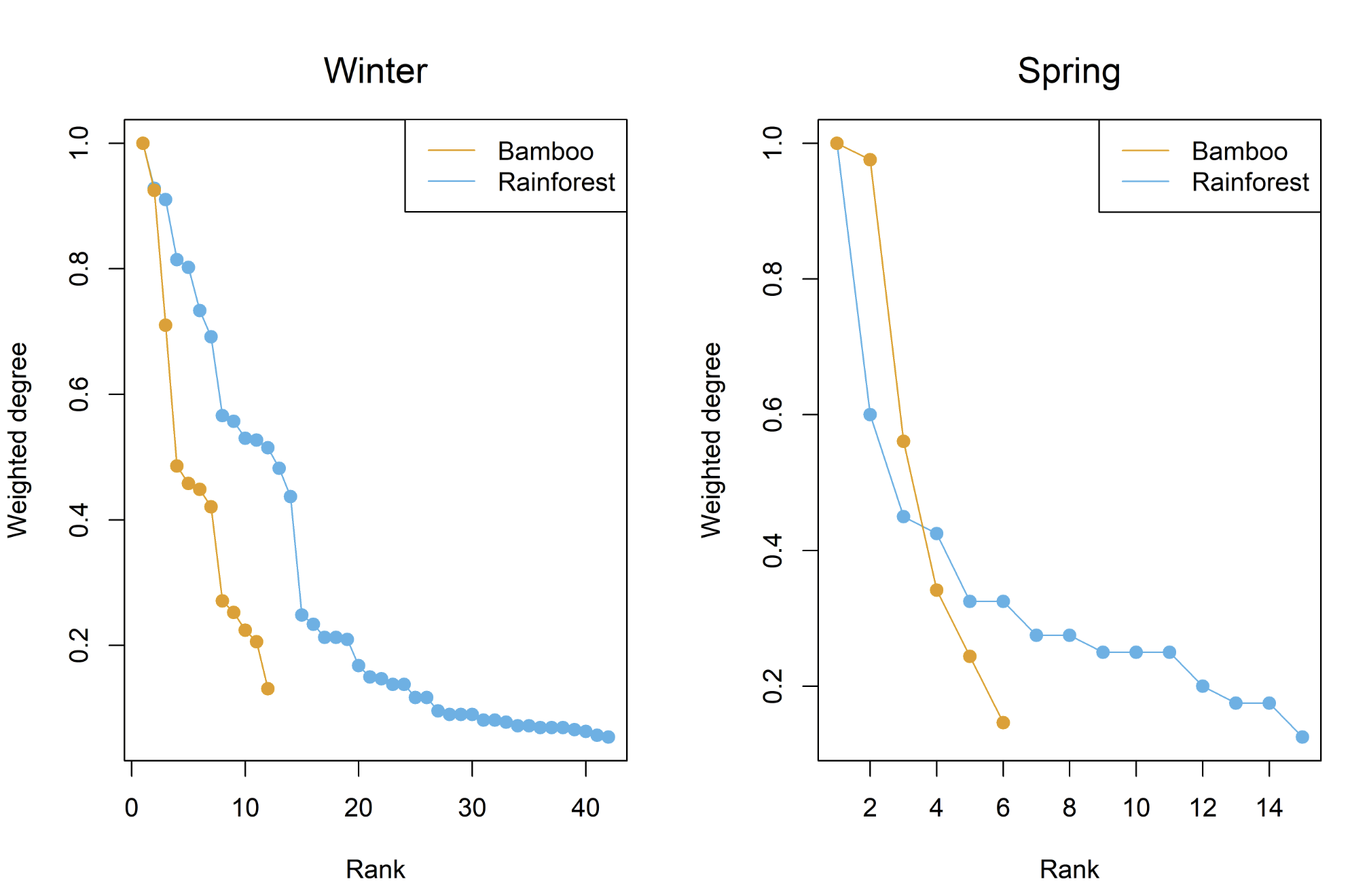

**Figure S2** - Normalised weighted degree distribution of flocks across seasons. The left panel represents flocks in winter and the right panel represents flocks in spring.

**TABLES**

**Table S1** - Definitions and biological interpretations of the network metrics used in this study. Adapted from (1).

| **Metric** | **Network/Node level** | **Definition** | **Biological Meaning** |
| --- | --- | --- | --- |
| Density | Network | The ratio of the number of actual connections out of all possible connections in the network | Gives information on the number of pair-wise interactions, out of all possible interactions in the flock. Higher values indicate a denser, highly connected network. |
| Modularity | Network | Calculates the strength of different modules/clusters of closely connected nodes in a network based on the number of edges within and across modules | Summarises how well-separated modules are in a network, and the aggregation of species to create distinct modules/subsets within a network. Higher values indicate when these subsets of species interact more strongly within them and weakly with the other species in the network. |
| Degree | Node | Number of edges connected to a particular node | Represents the number of interspecific associations of each species in a flock. |
| Weighted Degree | Node | Measures the frequency of distinct co-occurrences for every node | A centrality metric, represents the strength of interspecific associations for every species. Higher values indicate more importance of that species in the network. |
